# Phenotypic diversity of yeasts curated in the 5th edition of *The Yeasts*: A data-driven visualization approach

**DOI:** 10.64898/2026.02.15.706042

**Authors:** Taisuke Seike, Madoka Ide, Mari Yamamoto, Hiroya Yurimoto, Kosuke Shiraishi

**Affiliations:** Department of Bioscience and Bioinformatics, Graduate School Computer Science and Systems Engineering, Kyushu Institute of Technology, 680-4 Kawazu, Iizuka, Fukuoka 820-8502, Japan; Center for Biosystems Dynamics Research, RIKEN, 6-7-1 Minatojima-minamimachi, Chuo-ku, Kobe, Hyogo 650-0047, Japan; Department of Bioinformatics Engineering, Graduate School of Information Science and Technology, The University of Osaka, 1-5 Yamadaoka, Suita, Osaka 565-0871, Japan; Division of Applied Life Sciences, Graduate School of Agriculture, Kyoto University, Kitashirakawa-Oiwake, Sakyo-ku, Kyoto 606-8502, Japan

**Keywords:** The Yeasts, yeast phenotypes, phenotypic diversity, carbon utilization, fermentation capacity, data-driven analysis

## Abstract

Yeasts have long served as key experimental systems in genetics, cell biology, and fermentation research; however, these studies have largely focused on a limited number of model species. In contrast, yeasts are a fungal group with remarkable physiological and ecological diversity. To provide a global overview of yeast diversity, we compiled and visualized phenotypic information for approximately 1,300 yeast species documented in *The Yeasts* (5th edition), including carbon utilization profiles, fermentation capacity, growth temperature ranges, and reported isolation sources. Taxonomic reconciliation revealed extensive reannotation, with approximately 44% of the species undergoing name changes and recognized genera increasing from 143 to 233. Integrative analyses revealed pronounced phylogenetic structuring of metabolic breadth. Many basidiomycetous yeasts, particularly Agaricomycotina, exhibited broader, generalist-like carbon utilization, whereas ascomycetous yeasts, especially Saccharomycotina, more frequently displayed sugar-centered, specialist-like narrower profiles; model yeasts such as *Saccharomyces cerevisiae* and *Schizosaccharomyces pombe* fell at the narrow end of this spectrum. Fewer than half of all species fermented glucose, a trait largely confined to Saccharomycotina. In addition, nearly one-fifth of species failed to grow at 30 °C, the standard laboratory temperature. By reconstructing and visualizing decades of dispersed taxonomic knowledge accumulated in *The Yeasts*, this study reframes yeasts not merely as laboratory model organisms but as metabolically diverse fungi whose phenotypic diversity reflects diverse ecological contexts. The analytical framework presented here provides a foundation for integrating standardized quantitative phenotypes and newly described species and offers a starting point for exploring the latent ecological and metabolic potential of yeast diversity.

## 1 Introduction

Fungi constitute one of the most diverse kingdoms of life, encompassing organisms differing dramatically in their morphology, life cycle, and ecological strategies. Macroscopic fruiting body-forming fungi, filamentous molds, and unicellular yeasts are all members of this lineage, yet they exhibit fundamentally distinct modes of growth and reproduction (James et al., 2020). “Yeast” is not a taxonomic category but a growth form, typically referring to unicellular fungi (Dujon, 2010; Kurtzman et al., 2011; Nagy et al., 2014). Several fungal lineages exhibit dimorphic or polymorphic life cycles, switching between filamentous and yeast-like states in response to environmental conditions, and many basidiomycetes adopt a yeast-like form during part of their development (Bastidas & Heitman, 2009; Sánchez-Martínez & Pérez-Martín, 2001). Thus, yeasts represent a recurrent morphological and functional state rather than a single evolutionary lineage across the fungal kingdom (Floudas et al., 2012).

In laboratory, industrial, and educational contexts, the term “yeast” overwhelmingly refers to a few model species, most notably *Saccharomyces cerevisiae*, and to a lesser extent, *Schizosaccharomyces pombe* (Botstein & Fink, 2011; Hoffman et al., 2015). These species are widely associated with traits such as robust fermentation and growth at temperatures near 30 °C, shaping a canonical image of yeasts as sugar-fermenting microorganisms adapted to relatively warm environments (Hittinger et al., 2015; Kurtzman et al., 2011). However, this canonical image is derived from a narrow subsection of species and does not capture the full extent of the physiological and ecological variation found across fungal yeasts (Buzzini et al., 2017; Peter et al., 2018; Shen et al., 2018).

Yeasts exhibit striking heterogeneity in substrate utilization, fermentative capacity, thermal tolerance, and other physiological properties. Approximately 1,300 species are described in the 5th edition of *The Yeasts* (2011), and more than 2,000 species are currently recognized (Boekhout, 2021). Some non-model yeasts fail to grow at 30 °C, lack detectable fermentation, or utilize only a limited range of carbon substrates. These observations emphasize that the physiological characteristics of classical model yeasts are not universal across the fungal kingdom, highlighting the need for broader comparative perspectives.

A vast body of knowledge on yeast taxonomy, physiology, and ecology has been compiled over decades in *The Yeasts*, and its recent digital integration has accelerated the aggregation and accessibility of species-level information. However, most entries are organized as narrative descriptions or tabulated traits on a per-species basis. As a result, cross-species comparisons remain trivial, and it is difficult to intuitively grasp how phenotypes such as carbon substrate utilization, fermentation, and isolation sources, are distributed across yeast diversity. Despite the richness of the curated information, the absence of a comparative and visual framework limits our ability to recognize large-scale patterns or ecological biases, and the latent phenotypic potential of yeasts remains challenging to conceptualize.

To address this gap, we systematically curated and digitized phenotypic information for 1,262 species of yeasts described in the 5th edition of *The Yeasts*, and reorganized these data into a comparative and visualization-driven framework. Rather than generating new experimental data, the aim of this study was to reorganize existing literature-derived phenotypes, enabling cross-species comparisons and the identification of distributional biases. We placed particular emphasis on carbon substrate utilization and fermentative capacity, which are canonical traits often associated with yeasts, and examined their distributions at the phylum, subphylum, and family levels to extract features that are not apparent from species-by-species descriptions.

Our analyses revealed clear phylogenetic patterns in trait distributions, including contrasts between metabolic specialization and generalization, and highlighted the relationships between phenotypic traits and ecological contexts, such as isolation sources. Importantly, traits traditionally perceived as universal, such as fermentation or growth at 30 °C, were found to be far from representative of the yeast kingdom as a whole. Visualizing phenotypes as multidimensional distributions rather than isolated annotations provides a framework for reinterpreting yeasts not merely as laboratory or industrial organisms but also as diverse fungi adapted to a wide range of ecological niches.

By reorganizing literature-based phenotypic data into a comparative framework, this study provides a global overview of yeast diversity beyond species-by-species descriptions. The resulting patterns highlight lineage-associated trait distributions and prompt further quantitative and ecological investigation.

## 2 Materials and Methods

### 2.1 Data source and phenotypic curation

Phenotypic data were obtained from the 5th edition of *The Yeasts* (2011). In total, 1,262 yeast species listed in the volume were examined. Information regarding carbon and nitrogen substrate utilization, sugar fermentation, growth temperature, auxotrophy, genome GC content, major ubiquinone type, cell size, and reported isolation source was extracted (Table S1). Phenotypic data presented in tabular format were imported from PDF files and manually curated to correct transcription errors and inconsistencies. Traits described only in narrative text (e.g., cell size) were manually extracted. Carbon assimilation traits were recorded as reported in *The Yeasts*, and 34 substrates with relatively broad coverage across species were selected for comparative analyses. Fermentation ability was compiled for seven sugars (Table S1). Because the data were originally compiled for taxonomic descriptions rather than standardized physiological comparisons, assay conditions and scoring criteria varied across species. Only information contained in the 5th edition was used; subsequent taxonomic or phenotypic updates were not incorporated.

### 2.2 Taxonomic reconciliation

Because fungal nomenclature has undergone major revisions since the publication of the 5th edition, particularly following implementation of the “one fungus, one name” principle in the ICN (Turland et al., 2018), we reconciled species names using MycoBank (Robert et al., 2013) and updated nomenclature to valid names as of June 2025.

### 2.3 Retrieval of D1/D2 sequences and phylogenetic analysis

To infer genetic relationships among yeasts, we used sequences of the D1/D2 domain of 26S rDNA. Accession numbers provided in *The Yeasts* were used to retrieve sequences from NCBI. When multiple accession numbers were listed in the book, type strains were prioritized, and a representative sequence was selected when several type-derived entries existed. Sequences were aligned using MAFFT (default settings, v. 7.526) (Katoh & Standley, 2013), and phylogenetic trees were constructed using the neighbor-joining method implemented in MEGA 12 (Kumar et al., 2024) with the Kimura 2-parameter model (Kimura, 1980), pairwise deletion, and 1,000 bootstrap replicates. Species lacking D1/D2 sequences, possessing excessively short sequences unsuitable for alignment, or exhibiting unstable placement in preliminary analyses were excluded. In total, 1,192 out of the 1,238 species were included in the final tree.

### 2.4 Data processing and visualization

Phenotypic curation and visualization were performed using Python (v. 3.13.3). Carbon and nitrogen utilization traits were scored as binary or semi-quantitative values: positive = 1, weak/slow = 1, variable = 0.5, and negative = 0. For each species, the number of positively utilized substrates was computed and used for further analyses. Integration of phylogenetic information with phenotypic traits and generation of heatmaps and distribution plots were conducted using custom Python scripts.

### 2.5 Standardization of isolation source categories

Isolation sources in the *Ecology* section of *The Yeasts* are provided in a free-text format. To enable quantitative comparisons, relevant keywords and closely related terms were extracted from these descriptions and reclassified into ten operationally defined categories (categories 1–10). These categories included: plants, insects, other invertebrates, vertebrate animals, fungi and lichens, soil and terrestrial substrates, aquatic environments, air and artificial environments, food/fermentation/industrial environments, and unknown/unspecified sources. The classification criteria and associated keywords are presented in Table S2. The resulting category-level quantitative dataset is provided in Table S1. This categorization was not intended to define ecological niches, but instead to provide an operational framework to organize and compare reported isolation records across species.

### 2.6 Isolation source distribution analyses

Using the reclassified categories, the proportional composition of the isolated sources was calculated at both the phylum and subphylum levels. Comparative analyses were conducted between the two major fungal phyla (Ascomycota and Basidiomycota) and across the six yeast-forming subphyla represented in the dataset.

### 2.7 Geographic information and mapping of reported isolations

For each species, the reported countries of isolation were extracted from the *Ecology* section of *The Yeasts* and compiled by country. Each country was treated as an independent isolation record when a species was reported from multiple countries. The resulting country-level quantitative dataset is provided in Table S1. The compiled data were further organized into geographic information and visualized on a global map. It should be noted that these country-level data do not represent the natural distributions of yeasts but rather reflect the isolation records reported in the literature.

## 3 Results

### 3.1 Data collection and curation of phenotypic traits

We compiled literature-derived phenotypic annotations for 1,262 yeast species in the 5th edition of *The Yeasts* (Kurtzman et al., 2011), covering carbon assimilation (34 substrates), fermentation of seven sugars, growth across a temperature series, and additional descriptors including auxotrophy and isolation records (Table S1). Because these traits were originally collected for taxonomic descriptions, coverage and scoring depth varied across taxa, and some categories (e.g., nitrogen assimilation) contained substantial missingness. Accordingly, all downstream analyses were interpreted as comparative summaries of reported annotations rather than standardized quantitative measurements.

### 3.2 Taxonomic reconciliation after the publication of *The Yeasts*

A major challenge in integrating phenotypic data was the extensive revision of fungal taxonomy and nomenclature following the publication of *The Yeasts* (2011). The implementation of the International Code of Nomenclature for Algae, Fungi, and Plants (ICN) in 2013 introduced the “one fungus, one name” principle, abolishing the dual use of teleomorphic and anamorphic names. This update resulted in the large-scale reclassification of many asexual yeasts, including numerous former members of the genus *Candida*, on the basis of molecular phylogeny.

To account for these changes, all 1,262 species were reconciled against MycoBank and standardized to their currently accepted names as of June 2025. In total, 547 species (approximately 44%) underwent revisions in either genus or species names, and the number of genera increased from 143 to 233 (approximately 1.5-fold). Several taxa previously treated as distinct species were consolidated, resulting in a final dataset comprising 1,238 species belonging to 19 classes, 42 orders, 69 families, and 233 genera (Table S1). This taxonomic reconciliation enabled subsequent analyses of phenotypic distributions and isolation sources to be interpreted consistently within the current classification framework.

### 3.3 Phylogenetic framework of yeast diversity based on D1/D2 sequences

We reconstructed a phylogenetic framework for 1,238 yeast species listed in *The Yeasts* based on the sequences of the D1/D2 domain of 26S rDNA. D1/D2 sequences are widely used for yeast identification and classification, and their extensive use in previous taxonomic studies renders them suitable for comparative analyses (Kurtzman & Robnett, 1998; Kurtzman & Robnett, 2003). Sequences were retrieved from NCBI using the accession numbers provided in *The Yeasts*. When multiple accession numbers were listed in the book, a representative sequence was selected. The resolution of D1/D2 sequences has been reported to be insufficient for some yeast lineages (Daniel & Meyer, 2003; Lachance et al., 2003), and in our dataset, not all 1,238 species were consistently placed in stable tree topologies. The resulting D1/D2-based trees captured yeast-forming lineages across all six subphyla represented in the dataset and provided a practical phylogenetic scaffold for mapping phenotypic traits (Figure 1).

**Figure 1.**
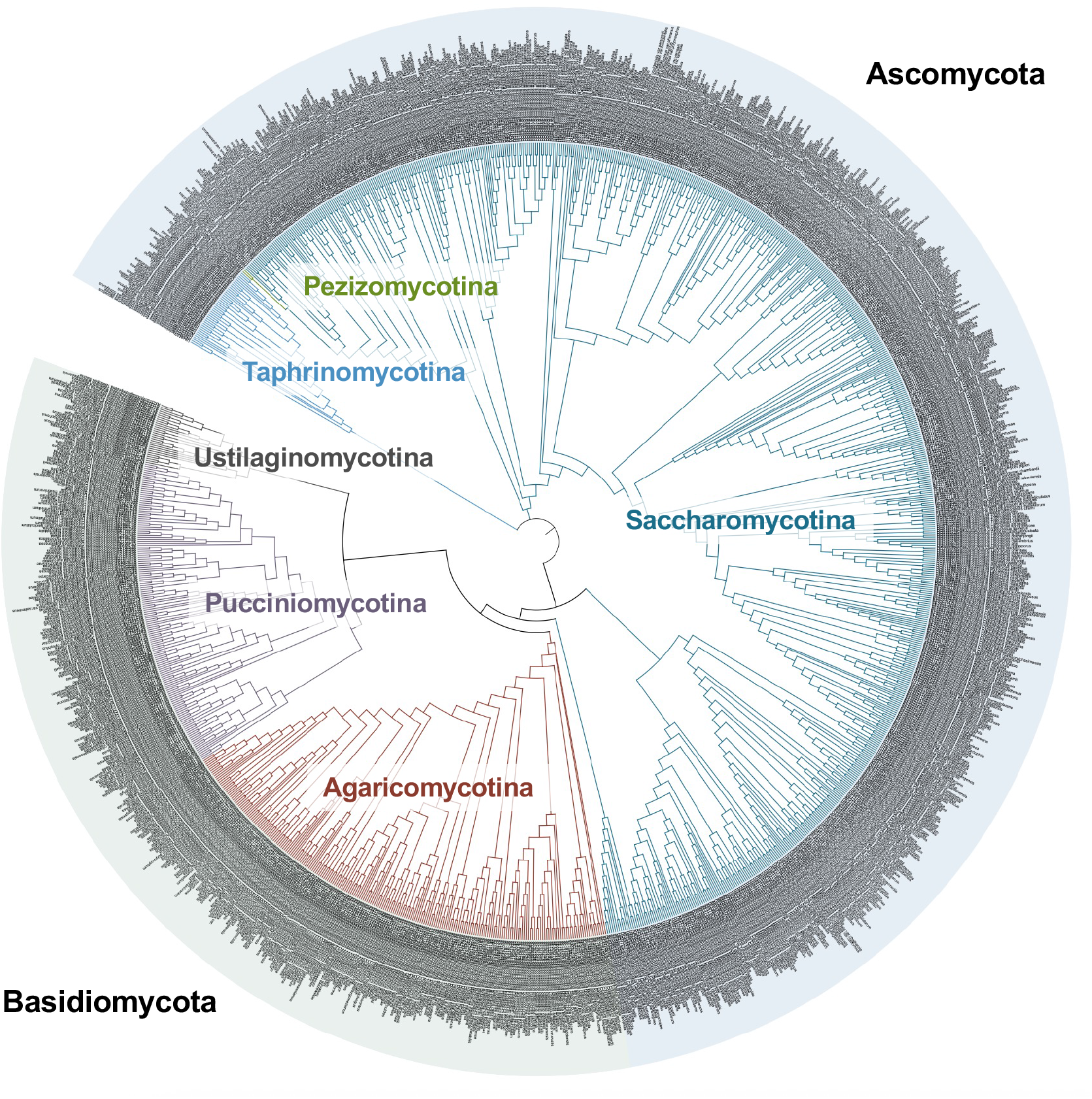
Phylogenetic framework of yeast diversity based on D1/D2 rDNA sequences. Phylogenetic tree of 1,192 yeast species inferred from D1/D2 domain sequences of 26S rDNA. The tree was constructed using the neighbor-joining method with the Kimura 2-parameter model and 1,000 bootstrap replicates. Branches are colored according to subphylum-level classification, representing the six fungal subphyla known to include yeast-forming species. Species belonging to Ascomycota and Basidiomycota are additionally indicated by distinct colors.

All six fungal subphyla, including yeast-forming species, were represented in the resulting tree. Yeasts were broadly divided into Ascomycota and Basidiomycota, comprising approximately 66% and 34% of the species, respectively. Within Ascomycota, Saccharomycotina was the largest group (770 species), followed by Taphrinomycotina (43 species) and Pezizomycotina (3 species). Within the Basidiomycota, Agaricomycotina (231 species), Pucciniomycotina (157 species), and Ustilaginomycotina (34 species) were identified. Across the dataset, more than 60% of species belonged to Saccharomycotina. With only a few exceptions, the D1/D2-based phylogenetic relationships were sufficiently resolved to provide an effective framework for subsequent analyses of phenotypic traits and isolation sources.

### 3.4 Global patterns of carbon and nitrogen utilization across yeasts

Next, we focused on carbon utilization profiles. Although the set of carbon sources examined varies across species in *The Yeasts*, 34 carbon sources were relatively well represented and were therefore used in our analyses. The number of utilizable carbon sources was calculated for each species. Positive, weak, and slow utilization were scored as 1, variable as 0.5, and negative as 0. The resulting values represent an operational index of the reported metabolic breadth, rather than a quantitative measure of metabolic capability.

Across the 1,238 species, the mean number of utilizable carbon sources was 17.6 ± 7.4 (Figure 2A). The maximum and minimum values were 32 and 1, respectively. Notably, the model yeasts *Saccharomyces cerevisiae* (8.5) and *Schizosaccharomyces pombe* (4.5) exhibited substantially narrower metabolic breadths than that of the overall dataset, indicating that commonly studied model organisms do not necessarily represent the full diversity of yeast metabolic capabilities.

**Figure 2.**
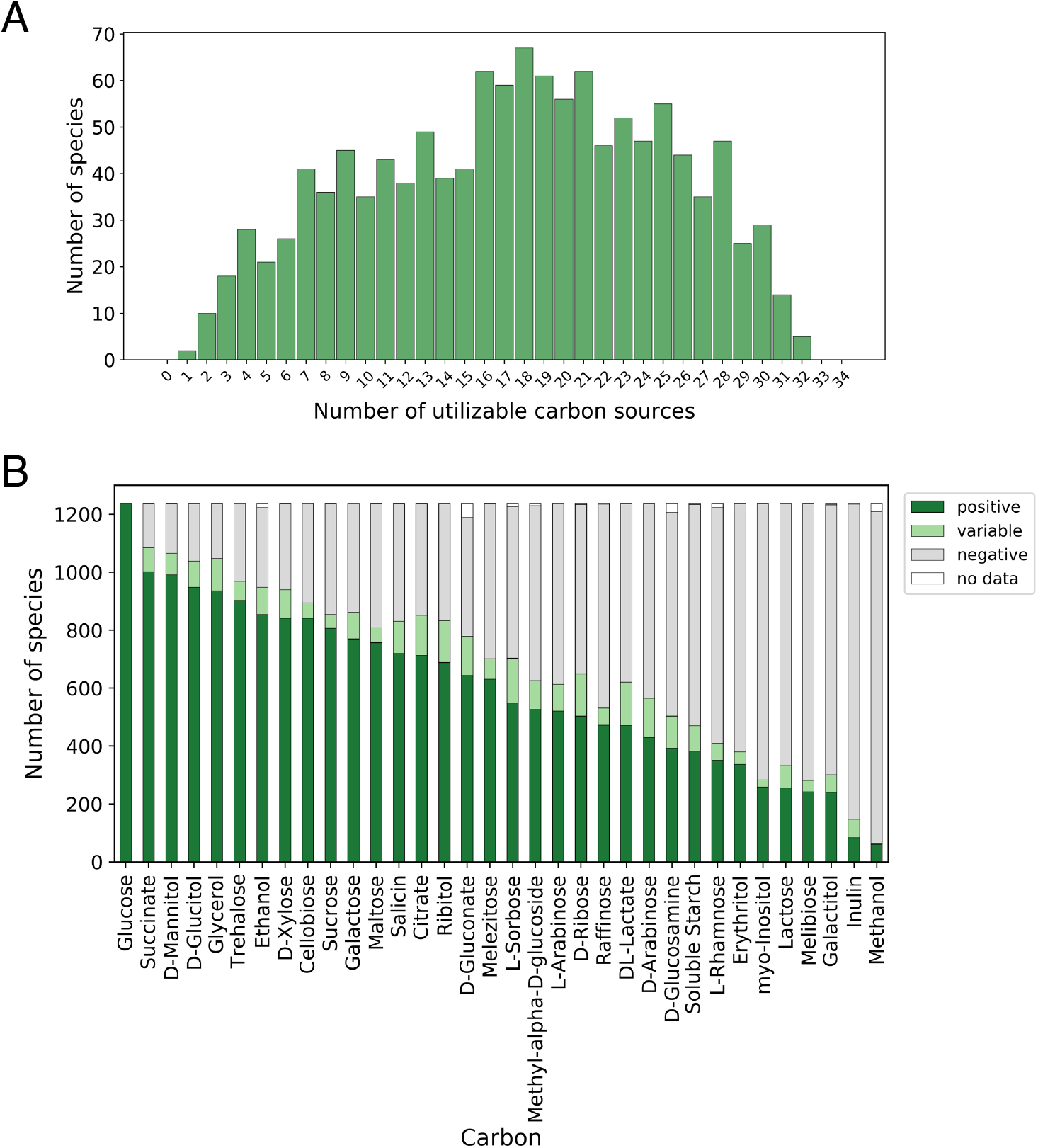
Global patterns of carbon substrate utilization across yeasts. (A) Distribution of carbon substrate utilization breadth across yeast species. For each species, the total number of utilizable carbon sources was calculated across 34 substrates based on data compiled from *The Yeasts*. Carbon utilization was scored as positive (1), variable (0.5), negative (0), or no data, as described in the Materials and Methods. The distribution of summed utilization scores across species (n = 1,238) is shown as a histogram, with the x-axis indicating the number of utilizable carbon sources and the y-axis indicating the number of species. (B) Frequency of carbon substrate utilization across yeast species. For each of the 34 carbon sources, the number of yeast species reported to utilize the substrate was counted. The x-axis indicates individual carbon sources, and the y-axis represents the number of species (out of 1,238) reported as positive, variable, negative, or no data for utilization.

The proportion of yeast species capable of utilizing each carbon source is shown in Figure 2B. Glucose was shown to be utilizable by all species and was the most universally available carbon source. Succinate and D-mannitol were also widely utilized by most species (84% and 83%, respectively). In contrast, methanol utilization was restricted to approximately 5% of the species, predominantly within *Komagataella* and certain *Candida* species. Polysaccharides, such as inulin, were also limited to a small number of species. Carbon sources such as fructose were excluded from the analyses because of their low discriminative power (Endoh et al., 2021).

Nitrogen utilization profiles, compiled from annotations for five nitrogen sources, showed similar patterns of heterogeneity, with a mean nitrogen utilization score of 1.64 ± 1.5 across species (Figure S1). Both *S. cerevisiae* and *S. pombe* scored 0; however, nitrogen-related data frequently lacked entries, and commonly used nitrogen sources such as ammonium sulfate and glutamine were not included in *The Yeasts*. Therefore, nitrogen utilization scores should be interpreted within the context of taxonomic descriptions rather than as comprehensive physiological assessments.

Visualization of the carbon utilization scores across the phylogenetic framework revealed marked phylogenetic skew (Figure S2). Many species within Ascomycota, particularly Saccharomycotina members, predominantly utilized sugars and exhibited relatively narrow metabolic breadths. In contrast, species within Basidiomycota, most notably members of Agaricomycotina, often utilized a wider array of carbon sources including alcohols, organic acids, and polysaccharides. These contrasting trends suggest that Ascomycota members display a more specialist-like profile, whereas many Basidiomycota members exhibit generalist-like metabolic strategies.

### 3.5 Fermentation capacity across yeast diversity

We next examined fermentation capacity across yeasts. Fermentation ability was recorded as the presence or absence of fermentation of seven sugars (glucose, galactose, sucrose, maltose, lactose, raffinose, and trehalose). Although yeasts are commonly regarded as “fermentative microorganisms,” a global survey of *The Yeasts* revealed a markedly different picture. Fewer than half of the species ferment glucose, the most basic carbon source (~600 species; Figure 3A), demonstrating that fermentation is not a universal property of yeasts. The ability to ferment lactose was particularly rare (<0.3%; four species), whereas other sugars exhibited varying frequencies of fermentative capacity across species. As fermentation ability in *The Yeasts* was generally assessed under aerobic conditions, these records do not necessarily reflect fermentative performance under conditions or quantitative aspects such as rate or yield.

**Figure 3.**
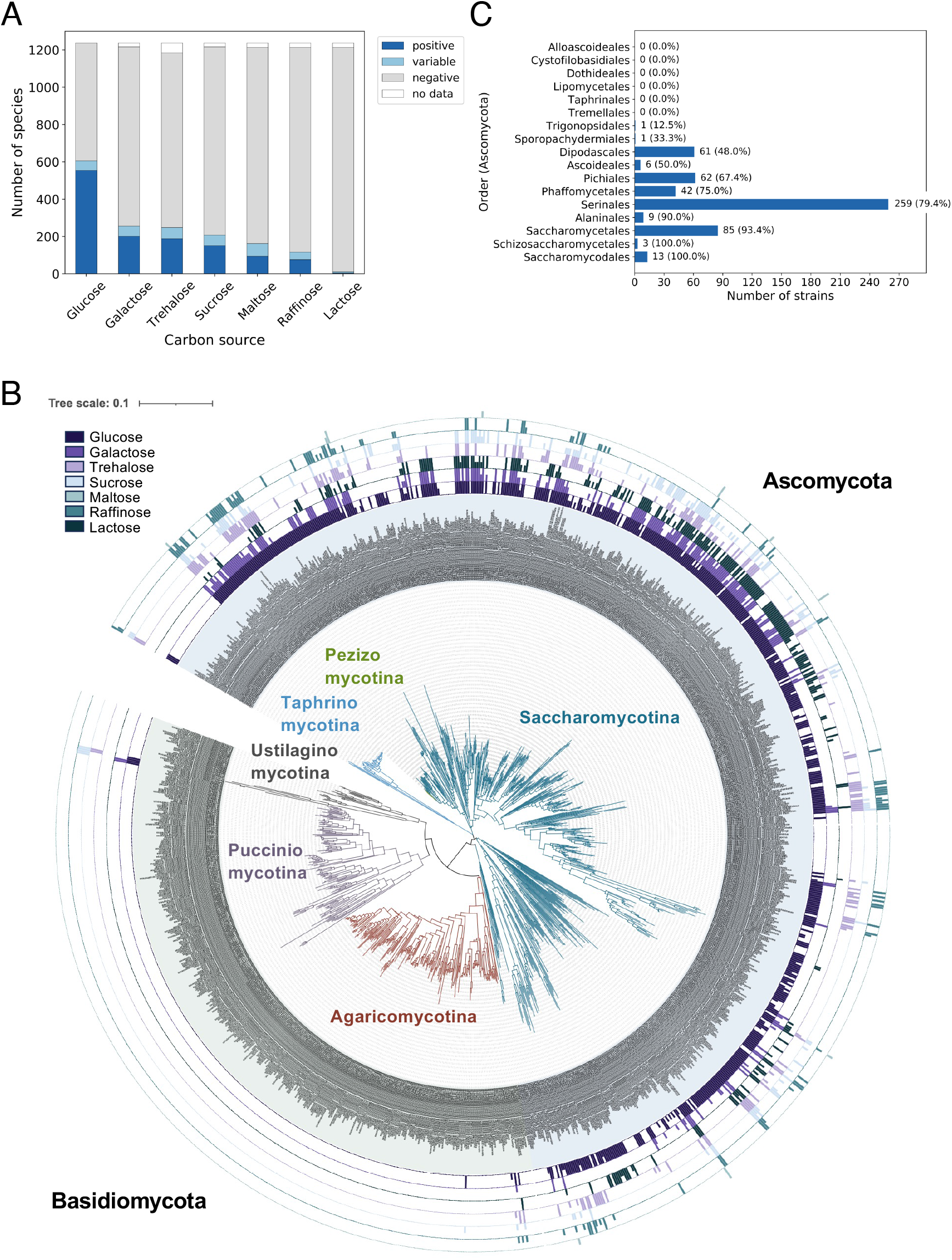
Fermentation capacity across yeast diversity. Distribution of sugar fermentation capacity across yeast species. For each species, fermentation ability was assessed for seven sugars (glucose, galactose, sucrose, maltose, lactose, raffinose, and trehalose) based on data compiled from *The Yeasts*. The x-axis represents individual species (n = 1,238), shown without labels due to scale, and the y-axis indicates the number of sugars reported as fermentable. Fermentation was scored as positive, variable, negative, or no data, as described in the Materials and Methods. Phylogenetic distribution of sugar fermentation capacity mapped onto the D1/D2-based yeast phylogeny. Fermentation abilities for the seven sugars (glucose, galactose, sucrose, maltose, lactose, raffinose, and trehalose) are shown as concentric bars surrounding the phylogenetic tree. Colored bars indicate the presence of fermentation capacity for each sugar. The tree was constructed using neighbor-joining based on D1/D2 sequences, and the scale bar represents 0.1 substitutions per site. (C) Proportion of glucose-fermenting species across the orders of Ascomycota. For each order within Ascomycota, the percentage of species reported to ferment glucose was calculated and visualized as a bar plot. The total number of species examined in each order and the corresponding percentage of glucose-fermenting species are indicated on the bars.

To investigate phylogenetic patterns, fermentation capacities across the seven sugars were mapped onto the D1/D2-based phylogeny (Figure 3B). Fermentation was extremely uncommon among Basidiomycota, where fermenting species were exceptional rather than representative. In contrast, fermenting species were more frequently observed within Ascomycota, particularly within Saccharomycotina. Notably, the presence of fermentative capacity in *Schizosaccharomyces* (Taphrinomycotina), including *S. pombe* and *S. japonicus*, is an exception to the overall phylogenetic pattern.

When Ascomycota were further examined at the order level, the orders were segregated into two broad groups: those containing fermentative species and those consisting almost exclusively of non-fermentative species (Figure 3C). These patterns indicate that fermentation is not a general feature of yeasts but instead has evolved in a phylogenetically restricted manner. The distribution of fermentation capacity partly corresponded to the carbon utilization profiles described above. Saccharomycotina frequently combine sugar-focused substrate utilization with fermentative capacity, whereas Basidiomycota tend to exhibit broad substrate utilization despite largely lacking fermentation. These differences may reflect ecological or metabolic strategies; however, quantitative comparisons were beyond the scope of the present dataset. More detailed analyses using standardized assays and quantitative performance metrics are required to further evaluate the evolutionary and ecological bases of these traits.

### 3.6 Growth temperature reveals unexpected limits of yeast diversity

Next, we examined the growth temperature profiles across yeasts. *The Yeasts* reports growth as presence/absence across multiple temperatures; here we curated and visualized data for 19, 25, 30, 35, 37, 40, and 45 °C (Figure 4). These records reflect qualitative assessments of growth and do not quantify growth rates or optima.

**Figure 4.**
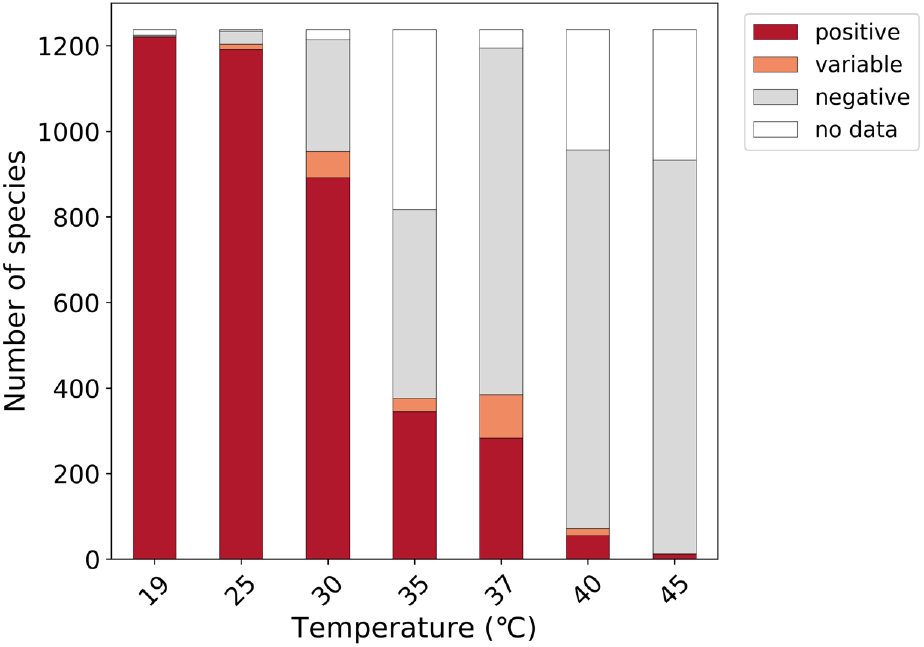
Growth temperature profiles across yeast species. Distribution of growth temperature tolerance across yeast species. For each species, growth ability was assessed at seven temperatures (19, 25, 30, 35, 37, 40, and 45 °C) on the basis of data compiled from *The Yeasts*. The x-axis represents individual species (n = 1,238), shown without labels due to scale, and the y-axis indicates the number of temperatures at which growth was reported. Growth was scored as positive, variable, negative, or no data, as described in the Materials and Methods.

Although 30 °C is widely used as the “standard” laboratory temperature for yeast cultivation, a global view of the dataset reveals that this convention is not universally representative. Approximately one-fifth of the species were unable to grow at 30 °C, whereas many grew at 25 °C or below. Therefore, commonly used cultivation temperatures may systematically exclude a sizeable fraction of yeast diversity. At the higher end of the temperature range, the number of species capable of growth declined sharply. Growth at 37 °C and 40 °C was restricted to a small subset of species (28% and 6.6%, respectively), and only 1.3% of yeasts grew at 45 °C, suggesting an upper limit of thermal tolerance near this range. These observations indicate that thermal adaptation in yeast is highly variable across species and differs markedly from canonical laboratory models.

Visualization of temperature-dependent growth highlights both the breadth and physiological limits of yeast diversity and underscores that “standard” laboratory conditions are not neutral with respect to diversity. Thus, temperature represents not only an experimental parameter but also a biological filter that shapes our view of yeast diversity and may influence isolation strategies in natural environments.

### 3.7 Distribution of the isolation sources

We organized information on yeast isolation sources reported in the *Ecology* section of *The Yeasts*, where environments or substrates are listed based on historical isolation records for each species. These records reflect documented isolation histories rather than natural biogeographic distributions and should be interpreted accordingly.

To enable comparative analyses, diverse free-text descriptions were reclassified into ten operational categories (Table S2). We first examined proportional distributions at the phylum level (Ascomycota vs. Basidiomycota; Figure 5). Isolates belonging to Ascomycota were more frequently associated with insects, whereas Basidiomycota were more often reported from plant- and soil-related sources. Subphylum-level comparisons revealed additional lineage-associated patterns (Figure 5; Figure S3). Insect-derived isolates were concentrated within Saccharomycotina, aquatic-associated records were enriched in Pucciniomycotina, and isolates from non-insect invertebrates were predominantly classified as Ustilaginomycotina. Projection of isolation source categories onto the D1/D2-based phylogeny further illustrated that these categories were partially clustered within particular lineages, consistent with the proportional summaries (Figure S4).

**Figure 5.**
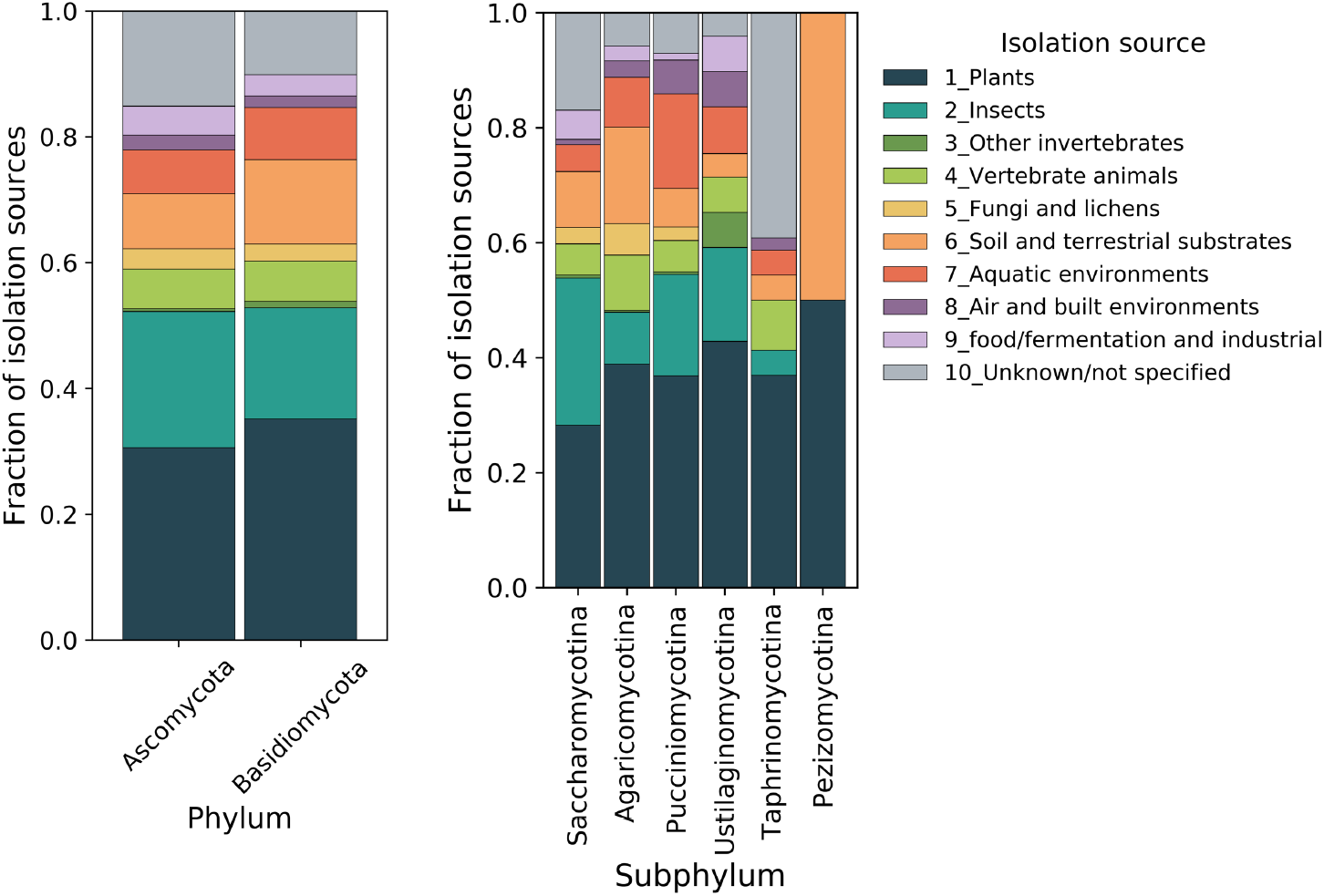
Taxonomic patterns of reported yeast isolation sources. Isolation source distributions were summarized at the phylum and subphylum levels. Isolation records were grouped into ten categories (categories 1–10; Table S2), and the proportional composition of each category was calculated within each taxonomic group. Stacked bars represent the fraction of isolation records assigned to each category.

Information on the isolation country was extracted for each species, compiled, and summarized (Figure S5). Isolation records were highly uneven across regions, with certain countries (e.g. the United States of America, Japan, and South Africa) accumulating disproportionately large numbers of reports, whereas many regions had few or no reports. These geographic biases more likely reflect differences in research history, fermentation and brewing cultures, and sampling effort, rather than true natural distributions. Mapping these data (Figure 6) further illustrates that the knowledge compiled in *The Yeasts* has been shaped by geographically concentrated research activities rather than by uniform global exploration.

**Figure 6.**
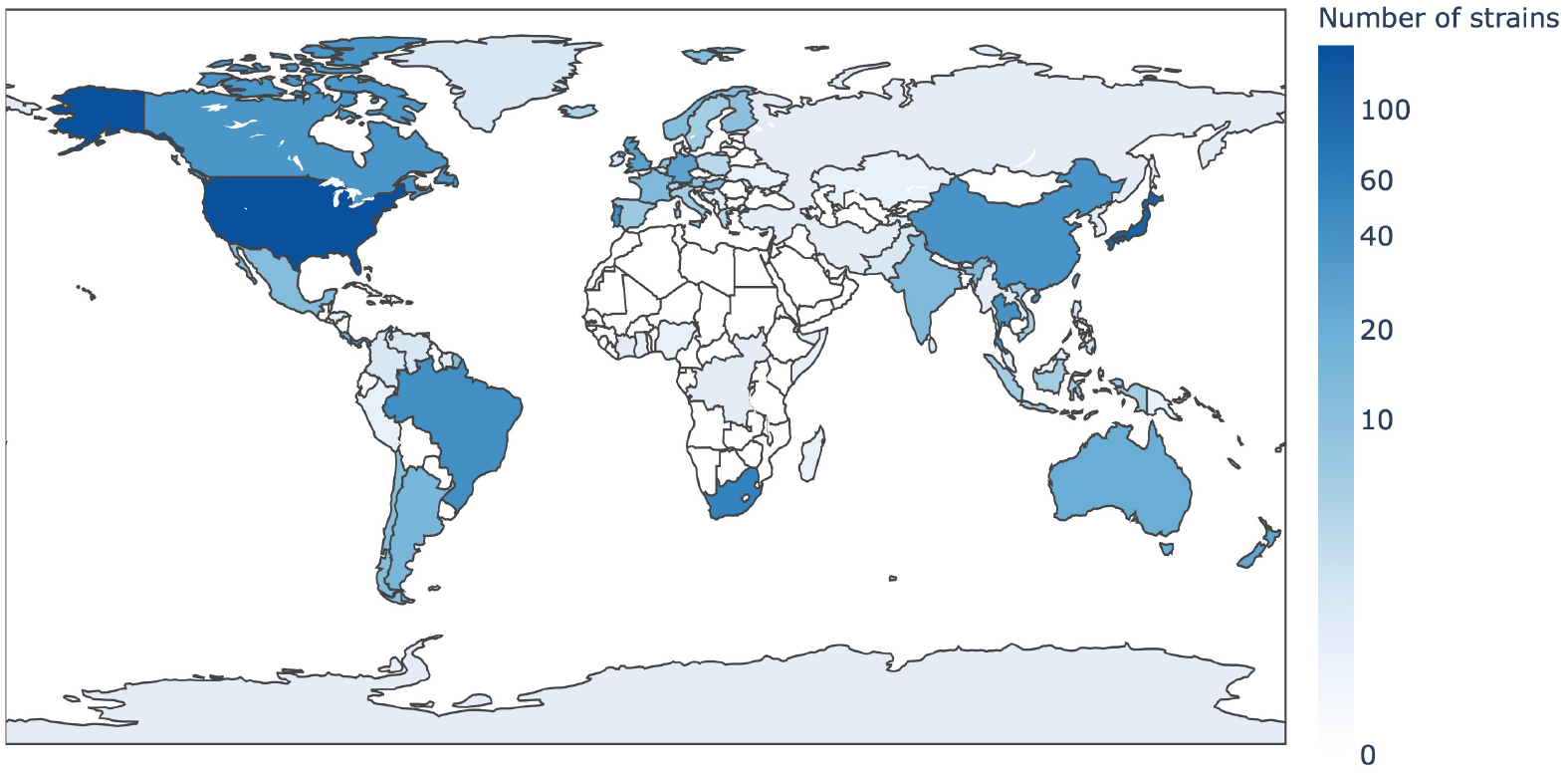
Global distribution of reported yeast isolation records. Country-level isolation information was extracted from the *Ecology* sections of *The Yeasts*. When a species was reported from multiple countries, each country was treated as an independent isolation record. Color intensity indicates the total number of reported isolation records per country. Color intensity indicates the total number of reported isolation records per country and reflects sampling effort rather than true species abundance.

By incorporating isolation sources into a global overview of yeast diversity, yeasts have emerged not simply as laboratory model organisms but as a diverse group adapted to a wide range of environments. Analyses of isolation records thus provide insights into yeast diversity and the historical processes through which yeasts have been explored and described, offering a perspective that reframes the conventional research-centric concept of “yeast.” These considerations highlight that phenotypes should be interpreted within ecological and historical contexts, which together shape our current view of yeast diversity.

## 4 Discussion

In this study, we systematically curated and visualized phenotypic information compiled in *The Yeasts* to provide a global overview of yeast diversity. By reorganizing literature-based data into a comparative framework, we offer a structured perspective on lineage-associated trait distributions beyond species-by-species descriptions. Our analyses revealed that fermentation capacity is not a universal feature among yeasts, but rather a trait that varies across phylogenetic lineages, consistent with a previous study of 40 yeast species (Hagman & Piškur, 2015). Likewise, growth temperature preferences showed considerable variability, with standard laboratory conditions (e.g. 30 °C) excluding a substantial fraction of species. Taken together, these results show that the conventional research- and industry-oriented view of “yeast” does not fully represent yeast diversity as a whole.

Yeasts are taxonomically widespread and ecologically heterogeneous (Liti et al., 2009) and exhibit considerable physiological diversity. Large-scale cross-species phenotypic summaries such as those presented here prompt testable hypotheses regarding yeast metabolic strategies. For example, the broader carbon utilization profiles frequently observed in many basidiomycetous yeasts may reflect greater metabolic flexibility under chemically diverse substrates, a possibility that can be evaluated using standardized growth assays. Similarly, the phylogenetically restricted distribution of glucose fermentation suggests lineage-associated constraints or repeated evolutionary transitions that warrant investigation through comparative genomic analyses. The visualizations presented here thus serve as an initial framework for exploring phenotypic breadth across yeast lineages.

This study has limitations inherent to the structure of the underlying data. The phenotypic information curated here is based on the 5th edition of *The Yeasts* and therefore excludes recently described species, particularly wild yeasts reported over the past decade. Moreover, the phenotypic data compiled in *The Yeasts* were originally obtained for taxonomic purposes rather than for standardized comparisons and thus differ in assay conditions, scoring criteria, and reporting depth. Accordingly, the distributions presented here represent interpreted summaries of reported annotations rather than quantitative measurements of physiological capacity.

Analyses incorporating isolation source information revealed lineage-associated biases within the curated dataset. Reported isolation records for Ascomycota frequently include insects and related substrates listed in the dataset, whereas Basidiomycota are more often documented from plant- and soil-related substrates. These contrasts are partly consistent with the phylogenetic biases observed in the carbon utilization profiles and may reflect a contrast between more generalized (generalist-like) and specialized (specialist-like) metabolic strategies. However, isolation records likely reflect research history, sampling efforts, food culture, and the development of fermentation industries (Boekhout et al., 2022; Harrison et al., 2024; Lachance, 2006), and therefore should not be interpreted as direct representations of natural ecological niches.

In addition, it is important to note that *The Yeasts* is no longer a static reference. It is currently maintained as a web-based database that is continuously updated. This online resource already includes phenotypic and ecological information for species that were not covered in the 5th edition, and it continues to expand with newly described taxa and additional trait descriptions. Consequently, the dataset analyzed in this study should be viewed as a snapshot derived from a specific edition, rather than as a complete representation of the currently available yeast diversity.

The existence of an actively curated and expandable database provides a clear opportunity for future extensions to the present framework. Incorporating newly added species, additional phenotypic traits not analyzed here, and more refined ecological annotations will enable increasingly comprehensive and up-to-date views of yeast diversity. Moreover, linking phenotypic information with genomic data, including gene content, metabolic pathways, and evolutionary relationships, is essential for advancing our understanding of the generation and maintenance of physiological diversity across yeast lineages.

To further advance our understanding of yeast diversity and metabolic potential, future efforts will benefit from integrating quantitative measurements obtained under standardized conditions, including carbon and nitrogen utilization; environmental tolerances such as temperature, pH, osmotic stress, and oxidative stress; growth rates; and fermentation yields. In parallel, a more refined organization of isolation source information will allow for a more precise assessment of how ecological contexts relate to phenotypic traits. The semi-structured nature of the current literature data also suggests that automated text extraction and AI-assisted data aggregation will facilitate a transition from resources that are primarily “read” to resources that can be actively “queried” for hypothesis generation.

By visualizing and reassembling the phenotypic information accumulated in *The Yeasts*, this study provides a new perspective for viewing yeast diversity on a global scale. Integrating information on carbon utilization, fermentation capacity, temperature tolerance, and isolation sources allows implicit assumptions about “standard” yeast traits and laboratory conditions to be reconsidered, revealing yeasts as a diverse group of organisms adapted to a wide range of environments. The framework presented here offers a neutral starting point for exploring yeast phenotypes and metabolic potential and provides a foundation for subsequent analyses incorporating standardized quantitative measurements and trade-off relationships.

## Author Contributions

Taisuke Seike conceived and designed the study, curated and analyzed the data, wrote the original manuscript, and reviewed and edited the manuscript for finalization. Madoka Ide and Mari Yamamoto curated and analyzed the data. Hiroya Yurimoto analyzed the data and reviewed and edited the manuscript. Kosuke Shiraishi conceived and designed the study, curated and analyzed the data, wrote the original manuscript, and reviewed and edited the manuscript. All the authors approved the final version of the manuscript.

## Acknowledgements

We thank Dr. Yuhei Goto for assistance with data curation and manual data entry. This work was supported by the GteX program (Grant Number JPMJGX23B4) (TS and KS).

## Data Availability Statement

The data supporting the findings of this study are available within the article and

Supplementary Information. Additional curated datasets generated in this study are available from the corresponding authors upon reasonable request.

## Supporting Information

Additional supporting information can be found online in the Supporting Information section.

## Legends to Supplementary Figures

**Figure S1 | Distribution of nitrogen source assimilation across yeast species**.

Distribution of nitrogen substrate utilization breadth across yeast species. For each species, the number of nitrogen sources reported as utilizable among five substrates was calculated based on data compiled from *The Yeasts*. The x-axis represents individual species (n = 1,238), shown without labels due to scale, and the y-axis indicates the number of utilizable nitrogen sources. Utilization was scored as positive, variable, negative, or no data, as described in the Materials and Methods.

**Figure S2 | Phylogenetic distribution of carbon substrate utilization breadth across D1/D2-based yeast phylogeny**.

The summed carbon substrate utilization score for each species, calculated as in Figure 2A (positive = 1, variable = 0.5, negative = 0), was mapped onto D1/D2-based yeast phylogeny. For each species, carbon utilization breadth is represented by the length of an adjacent bar, corresponding to the number of utilizable carbon sources. A vertical reference line indicates the midpoint of the distribution (17 substrates).

**Figure S3 | Isolation-source composition within each subphylum**.

Isolation records were grouped into ten operational categories (Table S2), and the proportional composition of each category was calculated within each yeast-forming subphylum. Rows are ordered by major phylum (Ascomycota followed by Basidiomycota). Cell annotations show the percentage contribution of each isolation-source category within a given subphylum.

**Figure S4 | Phylogenetic distribution of isolation source categories across the D1/D2-based yeast phylogeny**.

Isolation source categories were mapped onto the D1/D2-based phylogenetic tree constructed from 26S rDNA sequences. For each species, reported isolation sources extracted from the *Ecology* sections of *The Yeasts* were reclassified into ten operationally defined categories (categories 1–10; see Table S2) and visualized as colored tracks surrounding the phylogeny. Species without available isolation source information are indicated as unknown. The phylogenetic framework is identical to that shown in Figure 3B.

**Figure S5 | Country-level summary of reported yeast isolation records**.

Isolation records were compiled from country information provided in the *Ecology* sections of *The Yeasts*. Countries are ordered by decreasing number of reported isolations.

## Supplementary Tables

**Table S1** | **Curated phenotypic traits of 1**,**262 yeast species extracted from *The Yeasts* (5th edition)**.

**Table S2** | **Keywords and classification criteria used for isolation source categorization**.

