## Supplementary Figures for "Phenotypic diversity of yeasts curated in the 5th edition of *The Yeasts*: A data-driven visualization approach"

Fig. S1

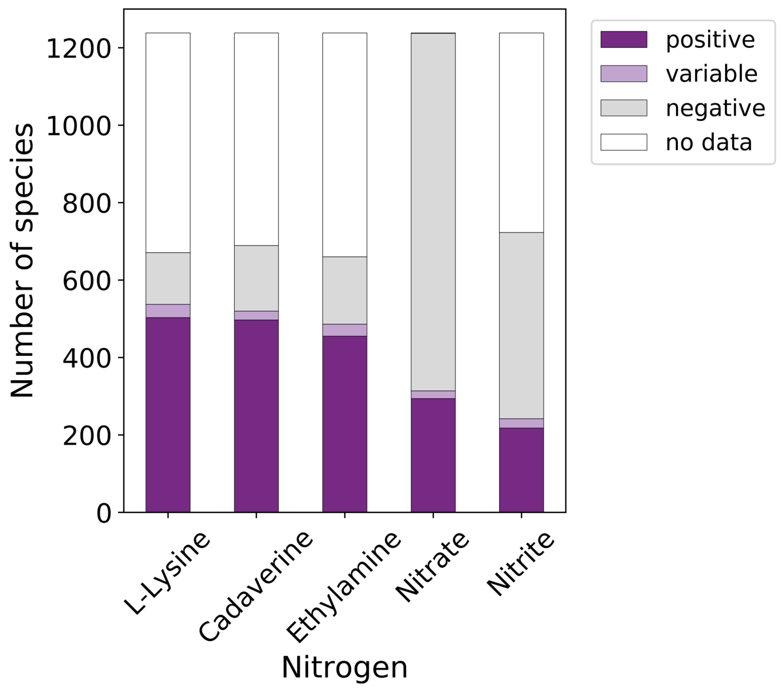


**Figure S1 | Distribution of nitrogen source assimilation across yeast species.**
Distribution of nitrogen substrate utilization breadth across yeast species. For each species, the number of nitrogen sources reported as utilizable among five substrates was calculated based on data compiled from *The Yeasts*. The x-axis represents individual species (n = 1,238), shown without labels due to scale, and the y-axis indicates the number of utilizable nitrogen sources. Utilization was scored as positive, variable, negative, or no data, as described in the Materials and Methods.

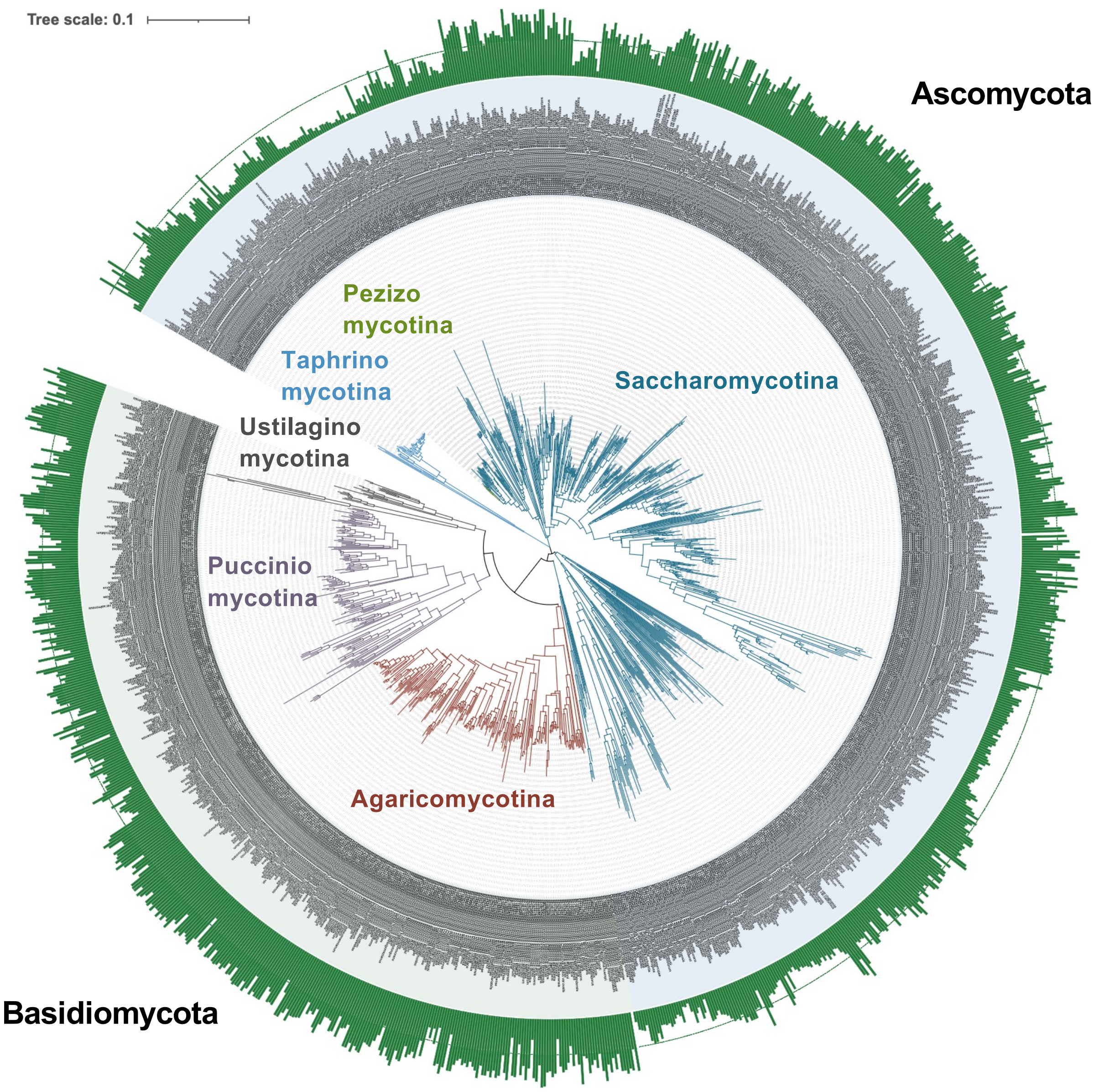


**Figure S2 | Phylogenetic distribution of carbon substrate utilization breadth across D1/D2-based yeast phylogeny.**
The summed carbon substrate utilization score for each species, calculated as in Figure 2A (positive = 1, variable = 0.5, negative = 0), was mapped onto D1/D2-based yeast phylogeny. For each species, carbon utilization breadth is represented by the length of an adjacent bar, corresponding to the number of utilizable carbon sources. A vertical reference line indicates the midpoint of the distribution (17 substrates).

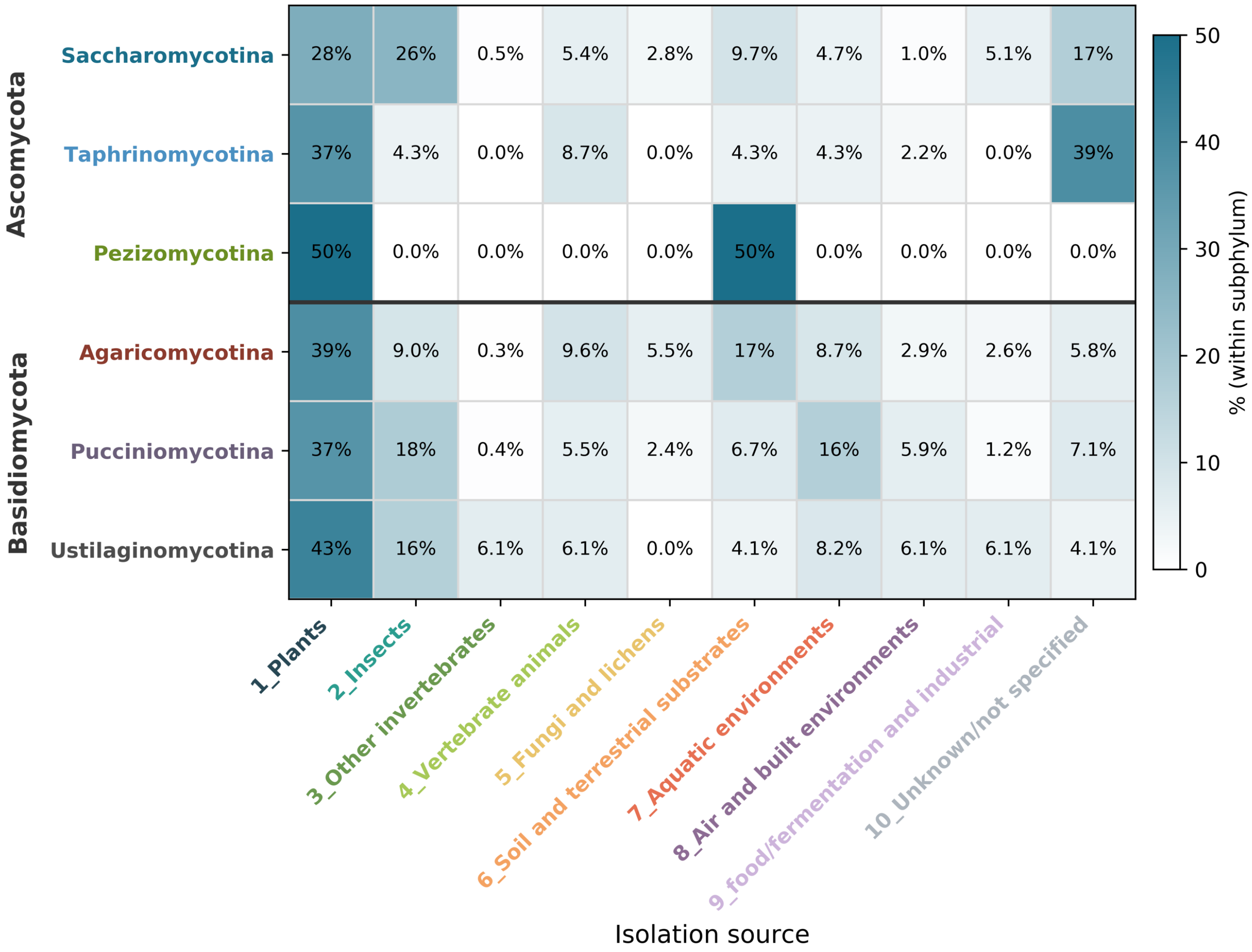


**Figure S3 | Isolation-source composition within each subphylum.**
Isolation records were grouped into ten operational categories (Table S2), and the proportional composition of each category was calculated within each yeast-forming subphylum. Rows are ordered by major phylum (Ascomycota followed by Basidiomycota). Cell annotations show the percentage contribution of each isolation-source category within a given subphylum.

Fig. S4

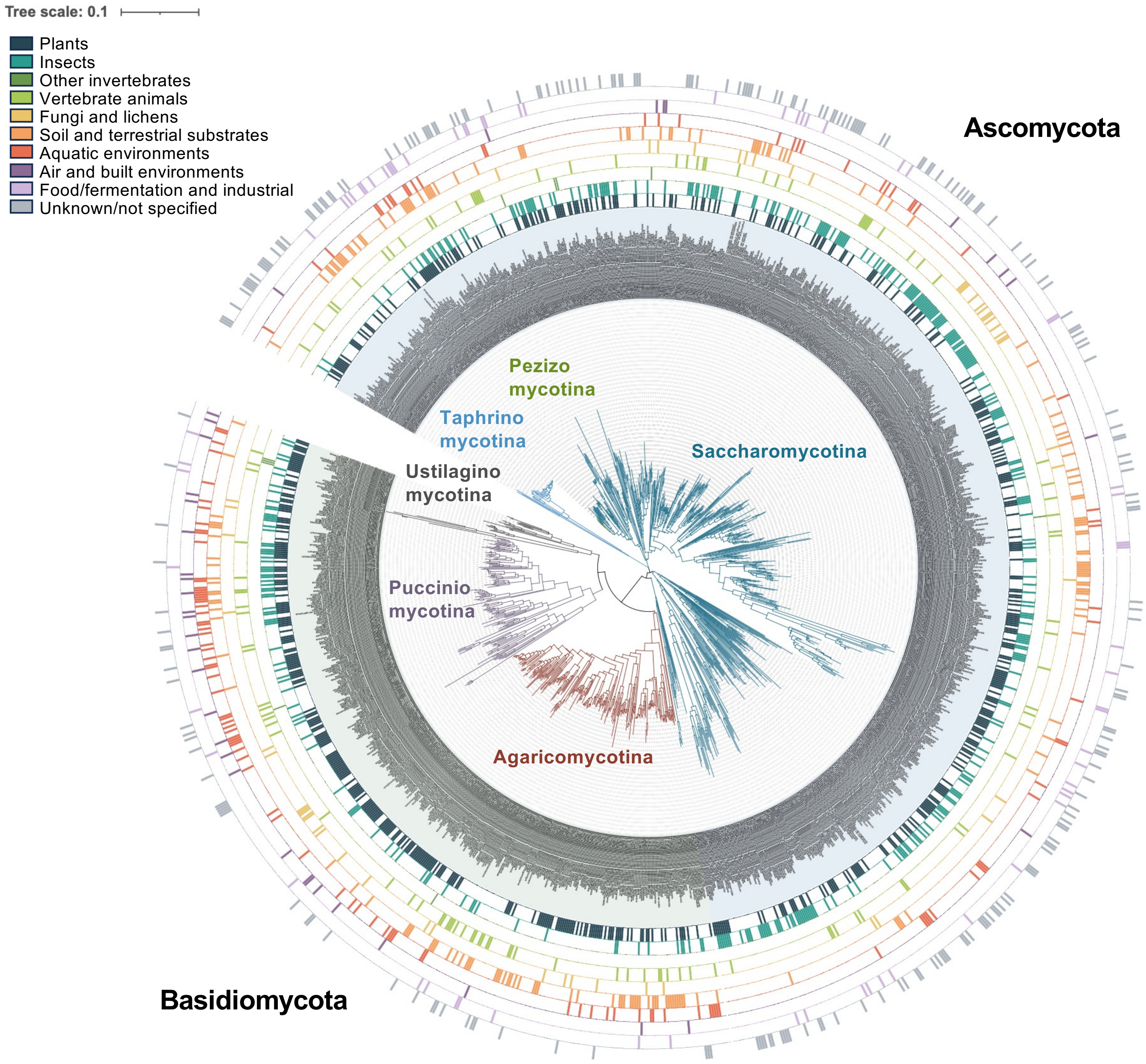


**Figure S4 | Phylogenetic distribution of isolation source categories across the D1/D2-based yeast phylogeny.**

Isolation source categories were mapped onto the D1/D2-based phylogenetic tree constructed from 26S rDNA sequences. For each species, reported isolation sources extracted from the Ecology sections of *The Yeasts* were reclassified into ten operationally defined categories (categories 1–10; see Table S2) and visualized as colored tracks surrounding the phylogeny. Species without available isolation source information are indicated as unknown. The phylogenetic framework is identical to that shown in Figure 3B.

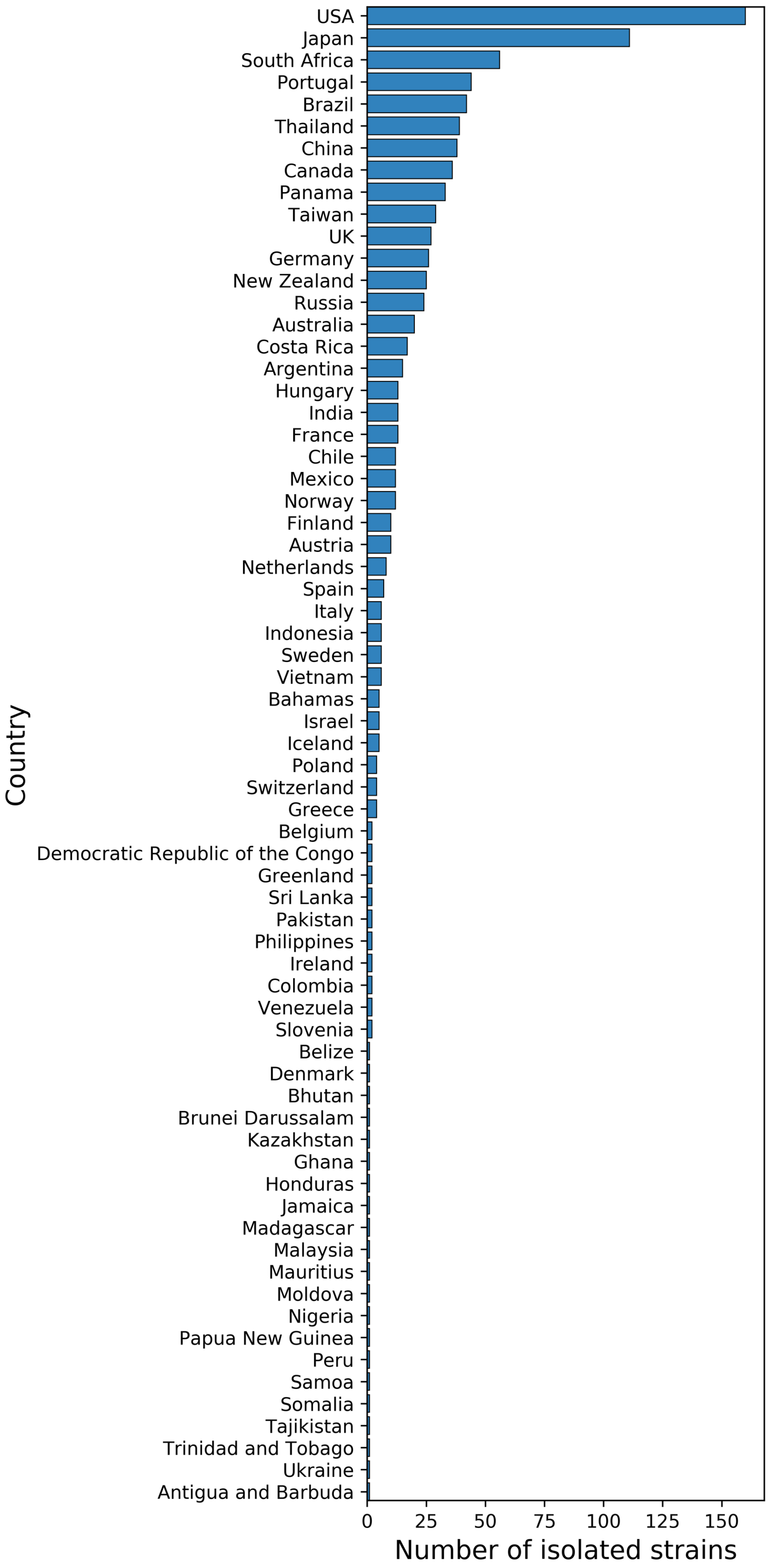


**Figure S5 | Country-level summary of reported yeast isolation records**
Isolation records were compiled from country information provided in the Ecology sections of *The Yeasts*. Countries are ordered by decreasing number of reported isolations.
